# Cohesin facilitates zygotic genome activation in zebrafishw

**DOI:** 10.1101/214023

**Authors:** Michael Meier, Jenny Grant, Amy Dowdle, Amarni Thomas, Jennifer Gerton, Philippe Collas, Justin M. O’Sullivan, Julia A. Horsfield

## Abstract

At zygotic genome activation (ZGA), changes in chromatin structure are associated with new transcription immediately following the maternal-to-zygotic transition (MZT). The nuclear architectural proteins, cohesin and CCCTC-binding factor (CTCF), contribute to chromatin structure and gene regulation. We show here that normal cohesin function is important for ZGA in zebrafish. Depletion of cohesin subunit Rad21 delays ZGA without affecting cell cycle progression. In contrast, CTCF depletion has little effect on ZGA whereas complete abrogation is lethal. Genome wide analysis of Rad21 binding reveals a change in distribution from pericentromeric satellite DNA, and few locations including the *miR-430* locus (whose products are responsible for maternal transcript degradation), to genes, as embryos progress through the MZT. After MZT, a subset of Rad21 binding occurs at genes dysregulated upon Rad21 depletion and overlaps pioneer factor Pou5f3, which activates early expressed genes. Rad21 depletion disrupts the formation of nucleoli and RNA polymerase II foci, suggestive of global defects in chromosome architecture. We propose that Rad21/cohesin redistribution to active areas of the genome is key to the establishment of chromosome organization and the embryonic developmental program.

**Author Summary:** During the first few hours of existence, early zygotic cellular events are regulated by maternally inherited molecules. From a defined timepoint, the zygotic genome gradually becomes active and is transcribed. How the zygotic genome is first held inactive before becoming rapidly activated is poorly understood. Both gene repression and activation mechanisms are involved, but one aspect that has not yet been investigated is how 3-dimensional chromosome structure influences genome activation. In this study, we used zebrafish embryos to model zygotic genome activation.

The multi-subunit protein complex, cohesin, and the DNA-binding protein CCCTC-binding factor (CTCF) both have well known and overlapping roles in 3-dimensional genome organization. We depleted cohesin subunit Rad21, or CTCF, to determine their effects on zygotic genome activation. Moderate Rad21 depletion delayed transition to zygotic gene expression, without disrupting the cell cycle. By contrast, moderate CTCF depletion had very little effect; however, strong depletion of CTCF was lethal. We surveyed genome-wide binding of Rad21 before and after the zygotic genome is activated, and determined what other chromatin factors and transcription factors coincide with Rad21 binding. Before genome activation, Rad21 was located at satellite DNA and a few noncoding genes, one of which (*miR-430*) is responsible for degrading maternal transcripts. Following genome activation, there was a mass relocation of Rad21 to genes, particularly active genes and those that are targets of transcriptional activators when the zygotic genome is switched on. Depletion of Rad21 also affected global chromosome structure.

Our study shows that cohesin binding redistributes to active RNA Polymerase II genes at the onset of zygotic gene transcription. Furthermore, we suggest that cohesin contributes to dynamic changes in chromosome architecture that occur upon zygotic genome activation.

## Introduction

Zygotic genome activation (ZGA) establishes, for the first time in a zygote, a genome that is competent for transcription (Blythe and Wieschaus, 2015; Fassnacht and Ciosk, 2017; Onichtchouk and Driever, 2016; Palfy et al., 2017; Svoboda et al., 2015). ZGA involves the transfer of maternal to zygotic control of embryonic development.

Commensurate with ZGA, maternal transcripts must be degraded at the maternal to zygotic transition (MZT) (Marco, 2017). In zebrafish, many maternal transcripts are targeted for degradation by *miR-430*, which is among the few early expressed transcripts in the embryo (Bazzini et al., 2012; Giraldez et al., 2006). Other maternal RNAs are N6-methyladenosine (m6A) modified, and are cleared by an m6A-binding protein, Ythdf2 (Zhao et al., 2017). Unique RNA-binding proteins also play a role in controlling RNA metabolism and turnover during MZT (Despic et al., 2017). Therefore, clearance of maternal RNA is essential for transition to the zygotic transcription program.

Mechanisms regulating both transcriptional activation and transcriptional repression are thought to control ZGA in the early embryo (Joseph et al., 2017; Lee et al., 2013; Onichtchouk and Driever, 2016). Evidence from Xenopus and zebrafish suggests the existence of a titratable, maternally deposited repressor that initially holds the transcription of the zygotic genome in check (Kimelman et al., 1987; Newport and Kirschner, 1982b; Nothias et al., 1995). In Xenopus, ZGA coincides with a dramatic increase in the nucleus-to-cytoplasm (N:C) volume ratio (Jevtic and Levy, 2015; Newport and Kirschner, 1982a). Increasing the N:C ratio by the addition of extra DNA (Newport and Kirschner, 1982b) or by injection of scaffolding proteins (Jevtic and Levy, 2015) accelerates ZGA. An *in vitro* study in Xenopus egg extracts showed that histones H3 and H4 are strong candidates for the maternal repressor activity (Amodeo et al., 2015); transcription repression by H3/H4 could be manipulated *in vitro* by altering the ratio of DNA template to histone quantities alone. In zebrafish, core histones outcompete transcription factors for access to the genome, thereby regulating the onset of transcription (Joseph et al., 2017). These studies suggest that transcription is activated as the histone repressors are titrated out during successive cell divisions. Therefore, up until ZGA, repression mechanisms counteract factors that activate transcription in the embryo.

Transcriptional activation at ZGA appears to involve a combination of ‘pioneer’ transcription factor activity, and a gain in active chromatin modifications. At the zebrafish maternal-to-zygotic transition (MZT), distinctive histone modifications appear (Andersen et al., 2013; Vastenhouw and Schier, 2012; Vastenhouw et al., 2010), and nucleosomes become strongly positioned at promoters (Zhang et al., 2014b). Even before ZGA, the zebrafish genome is marked with modified histones (Lindeman et al., 2011) and specific sites of DNA methylation (Jiang et al., 2013; Potok et al., 2013). However, although chromatin modifications can demarcate active regions of transcription, additional factors are usually needed for transcription activation (Hontelez et al., 2015). Sequence-specific transcription factors operating at ZGA vary between species. In Drosophila, the zinc finger protein, Zelda, activates many early genes (Harrison et al., 2011; Li et al., 2014). In zebrafish, Nanog, Pou5f3 (also called Oct4) and SoxB1 regulate expression of early zygotic genes (Lee et al., 2013; Leichsenring et al., 2013; Onichtchouk and Driever, 2016). Recently, the DUX family of transcription factors was found to activate zygotic genes in mice (De Iaco et al., 2017; Hendrickson et al., 2017; Whiddon et al., 2017).

Global chromatin structure is also linked to transcription activation; for example, formation of architectural features such as topologically associated domains (TADs) mark the onset of transcription in the mouse embryo (Flyamer et al., 2017; Lu et al., 2016). In Drosophila, chromatin architecture in the form of TADs emerges at ZGA independently of gene transcription (Hug et al., 2017). This is consistent with the idea that genome structure formation precedes transcription (Krijger and de Laat, 2017). Chromatin structure in turn influences the binding of transcription factors and RNA Polymerase II (RNAPII) (Newman and Young, 2010).

While individual players in ZGA may vary between species, a universal theme is that the spatial organization of chromosomes changes as cells commit to developmental fates (de Wit et al., 2013; Hug et al., 2017; Phillips-Cremins, 2014; Vietri Rudan et al., 2015). Spatial organization of the genome depends in part on the nuclear architectural proteins, cohesin and CCCTC-binding factor (CTCF), which contribute to the 3-dimensional (3D) organization of chromosomes (Vietri Rudan and Hadjur, 2015), and the formation of DNA loops within TADs (Giorgetti et al., 2014; Hug et al., 2017; Van Bortle et al., 2014; Vietri Rudan et al., 2015). Compartmentalization of active and inactive regions of the genome does not depend on CTCF (Nora et al., 2017) or cohesin (Merkenschlager and Nora, 2016). However, local spatial organization within TADs can facilitate transcription of developmental loci (Ferraiuolo et al., 2010; Narendra et al., 2015; Rousseau et al., 2014), thereby determining cell fate and driving embryo development.

In this study, we asked whether cohesin and CTCF contribute to ZGA. We found that cohesin (but not CTCF) depletion delays ZGA, and that chromosome bound cohesin spreads from satellite and non-coding DNA to genes when the zygotic genome becomes activated. A fraction of gene-associated cohesin binding sites are co-occupied by ‘pioneer’ transcription factors Pou5f3 and Sox2, and enriched for active histone marks. We propose that cohesin plays a crucial role in organizing a chromatin structure that is permissive for transcription at ZGA.

## Results

### Depletion of cohesin and CTCF in zebrafish embryos

As embryos progress through MZT at 3.3 hours post-fertilization (hpf), the main wave of zygotic gene transcription is activated (Fig. 1A, Heyn et al. (2014)). Relatively low levels of Rad21 (~100) and CTCF (~150) transcripts are present pre-ZGA, with both transcript and protein levels increasing by 2-3-fold in the main wave of ZGA (Fig. 1B,C). Genes encoding Rad21 and CTCF are essential for cell survival (Nasmyth and Haering, 2005) (including germ cells), which limits the genetic tools available for their manipulation. We previously bypassed homozygous lethality by using morpholino oligonucleotides (MO) to tightly titrate the levels of Rad21 and CTCF (Marsman et al., 2014; Rhodes et al., 2010; Schuster et al., 2015). Here, we were able to substantially reduce the protein levels of cohesin subunit Rad21 and CTCF in early embryos in order to assess their effects on zygotic genome activation (Figs 1, S1, S1E). Rad21-depleted embryos were rescued by a transcript encoding wild type Rad21, but not by mutant Rad21 containing the *rad21*^*nz171*^ nonsense mutation (Horsfield et al., 2007) (Fig. S2).

**Figure 1:**
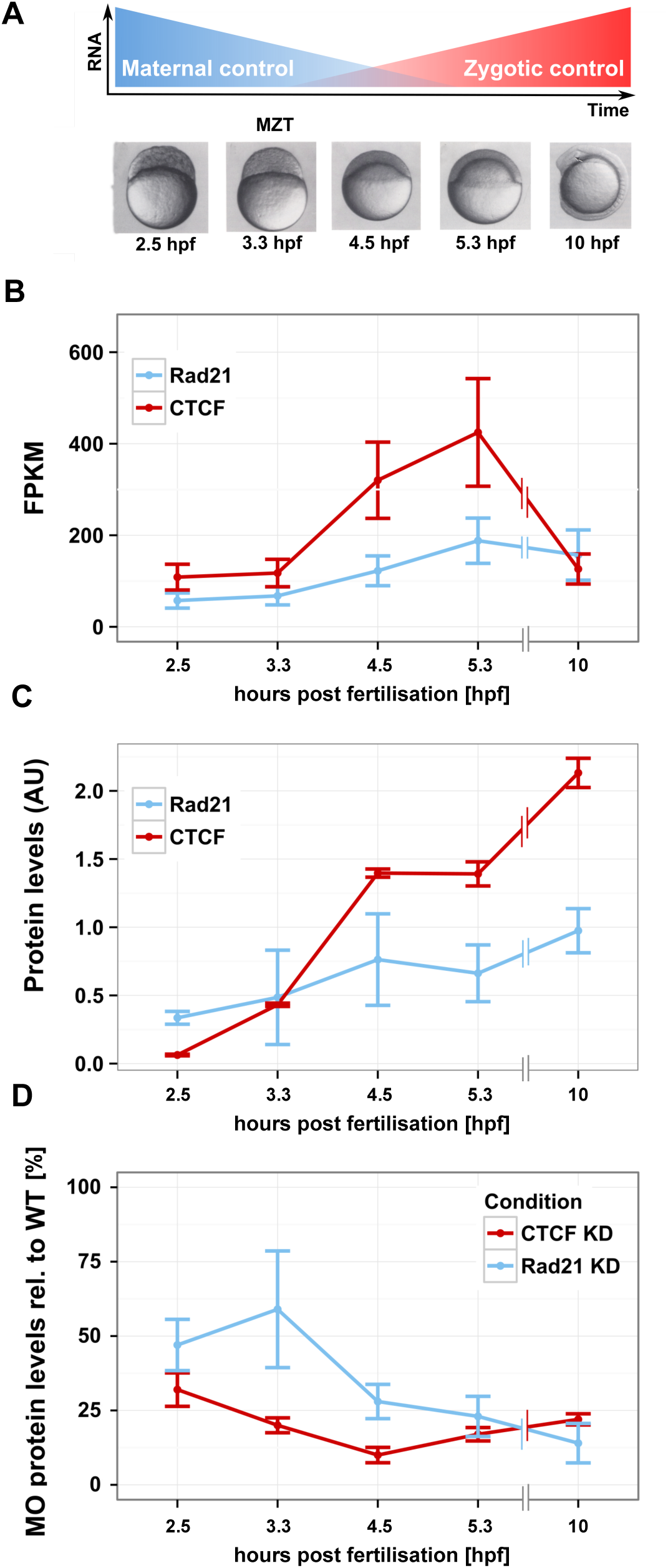
Rad21 and CTCF are present pre-ZGA and can be effectively depleted in early zebrafish development. (A) As embryos reach MZT, maternal transcripts are degraded and zygotic transcripts accumulate. The series of embryos below represents time points that were sampled for RNA-seq. hpf = hours post-fertilization. (B) Transcript numbers expressed as fragments per kilobase mapped (FPKM) of Rad21 and CTCF as measured by RNA-seq across the indicated time points. Error bars represent 95% confidence intervals. Quantitation of immunoblots for Rad21 and CTCF protein levels, normalized against those of γ-tubulin. Data are means ± s.d. n = 3. (D) Quantitation of immunoblots for Rad21 and CTCF protein levels, following depletion of these proteins using morpholino oligonucleotides (Rad21 knockdown (KD) and CTCF KD). Protein levels are expressed as a percent of wild type levels and were normalized against those of γ-tubulin. Images of all immunoblots are provided in Figs S1, S1E

MOs injected at the 1-cell stage reduced protein levels of Rad21 and CTCF by 40-80%, even pre-ZGA (Figs 1D, S1). By the 4.5 hpf ‘dome’ stage, Rad21-depleted embryos had just slightly fewer cells than wild type, although the difference was not significant (*p* = 0.1138, unpaired *t*-test) (Fig. S3A). Analysis of cell cycle status by flow cytometry showed that both wild-type and Rad21-depleted embryos had a large majority of cells with 2N DNA content, although the Rad21-depleted embryos had an increase in cells with 4N DNA content, from 7% in wildtype to 10% with Rad21 depletion (Fig. S3B). Surviving CTCF-depleted embryos displayed similar cell cycle profiles to wild type (data not shown). However, many CTCF MO-injected embryos died pre-MZT, suggesting that survivors had sub-threshold CTCF depletion.

### Rad21 depletion delays the onset of the zygotic transcription program

To determine the effects Rad21 and CTCF depletion on zygotic transcription, we used RNA-seq to analyse the transcriptome of untreated embryos (referred to here as ‘wild type’) and embryos treated with Rad21- or CTCF-targeting MOs. Five developmental stages were analyzed spanning pre-MZT (2.5 hpf), MZT (3.3 hpf), and post-MZT (4.5 and 5.3 hpf) stages up to the tailbud stage (10 hpf). The sample-to-sample distances between the expression profiles were calculated (R package DESeq2) to cluster time points and treatments. A graphical representation of the sample-to-sample distance is shown in a principal component (PCA) plot in Figure 2A. We found that Rad21 depletion results in a complement of transcripts that appear delayed in developmental timing relative to wild type at the post-MZT stages of 4.5 and 5.3 hpf (PC1, Fig. 2A). By contrast, profiles from CTCF-depleted embryos cluster similarly to wild type embryos from the same stage (Fig. 2A).

**Figure 2:**
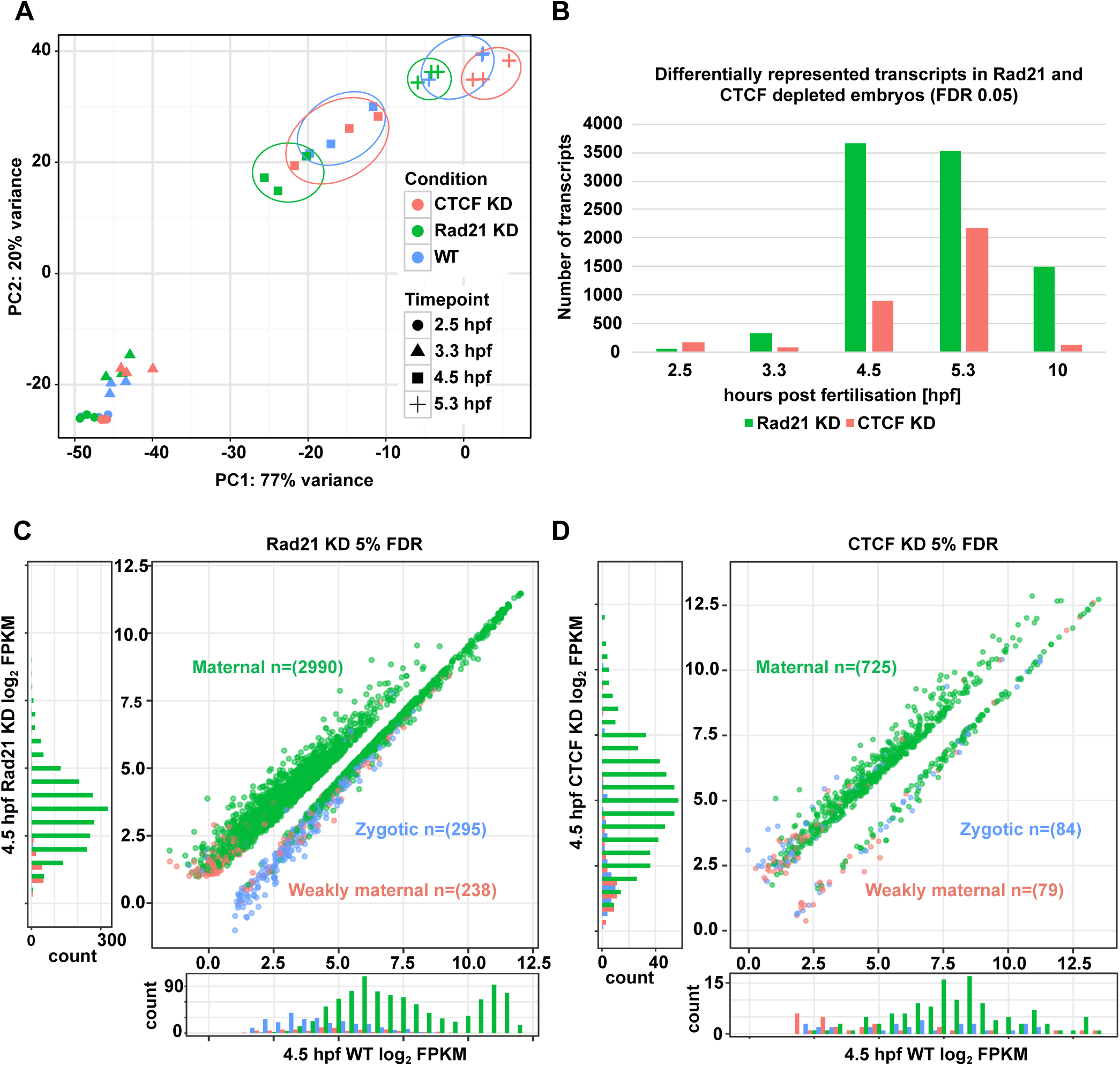
Rad21 depletion delays the onset of the zygotic transcription program. (A) PCA plot of RNA-seq triplicate samples for pools (n=100) of wild type (WT), Rad21-depleted (KD) and CTCF-depleted (KD) conditions at time points 2.5-5.3 hpf. PC1 and PC2, together accounting for 97% of the variation, identify sample separation by developmental time. Samples from different conditions (WT, Rad21 KD and CTCF KD) show clustered differences at 4.5 hpf and 5.3 hpf. (B) Number of differentially represented transcripts in Rad21 KD and CTCF KD embryos at stages 2.5-10 hpf. (C) Scatterplot of differentially represented transcripts (total = 3,253, FDR=0.05) between WT and Rad21 KD at 4.5 hpf. (D) Scatterplot of differentially represented transcripts (total = 888, FDR=0.05) between WT and CTCF KD at 4.5 hpf. Histograms depict the number of over-represented (y-axis) and under-represented transcripts (x-axis) in (C) and (D).

### More transcripts are affected following depletion of Rad21 than of CTCF

We identified differentially represented transcripts between wild type, Rad21- and CTCF-depleted embryos at each of the five stages (Table S1). Rad21 and CTCF depletion most robustly affected transcript levels post ZGA at 4.5 hpf and 5.3 hpf (Fig. 2B). We next annotated (Lee et al., 2013) the origin of the differentially represented transcripts (maternal, weakly maternal or zygotic) in Rad21-depleted embryos and CTCF-depleted embryos compared to wild type at the 4.5 hpf ‘dome stage’. We found that upon Rad21 depletion, 3,285 differentially represented transcripts (FDR=0.05) were both maternal and zygotic, with maternal transcripts more abundant relative to wild type and zygotic transcripts under-represented (Fig. 2C, Table S1), suggesting ZGA is delayed. Following CTCF depletion, there were 888 differentially represented transcripts, almost 4-fold fewer than observed upon Rad21 depletion (FDR=0.05) (Fig. 2D, Table S1).

We then plotted the expression levels of differentially expressed transcripts identified from dome stage at all time points sampled. Transcripts that are under-represented in Rad21-depleted embryos normally increase over developmental time in wild type, and transcripts that are over-represented upon Rad21 depletion are reduced over time in wild type embryos (Fig. 3A,B), with significantly more transcripts affected when compared to CTCF depletion (Fig. S4). CTCF-depleted embryos showed a similar trajectory of differentially represented transcripts over developmental time (Fig. S4A,B). Following Rad21 depletion, delay in the expression of individual zygotic genes was confirmed by quantitative PCR (Fig. 3C). Furthermore, expression of these genes was similarly delayed by depletion of a second cohesin subunit, Smc3 (Fig. 3C, Fig. S1E), suggesting that the effect of Rad21 depletion on ZGA is mediated through abolition of cohesin complex function. We conclude that even partial depletion of cohesin causes a delay in zygotic genome activation.

**Figure 3:**
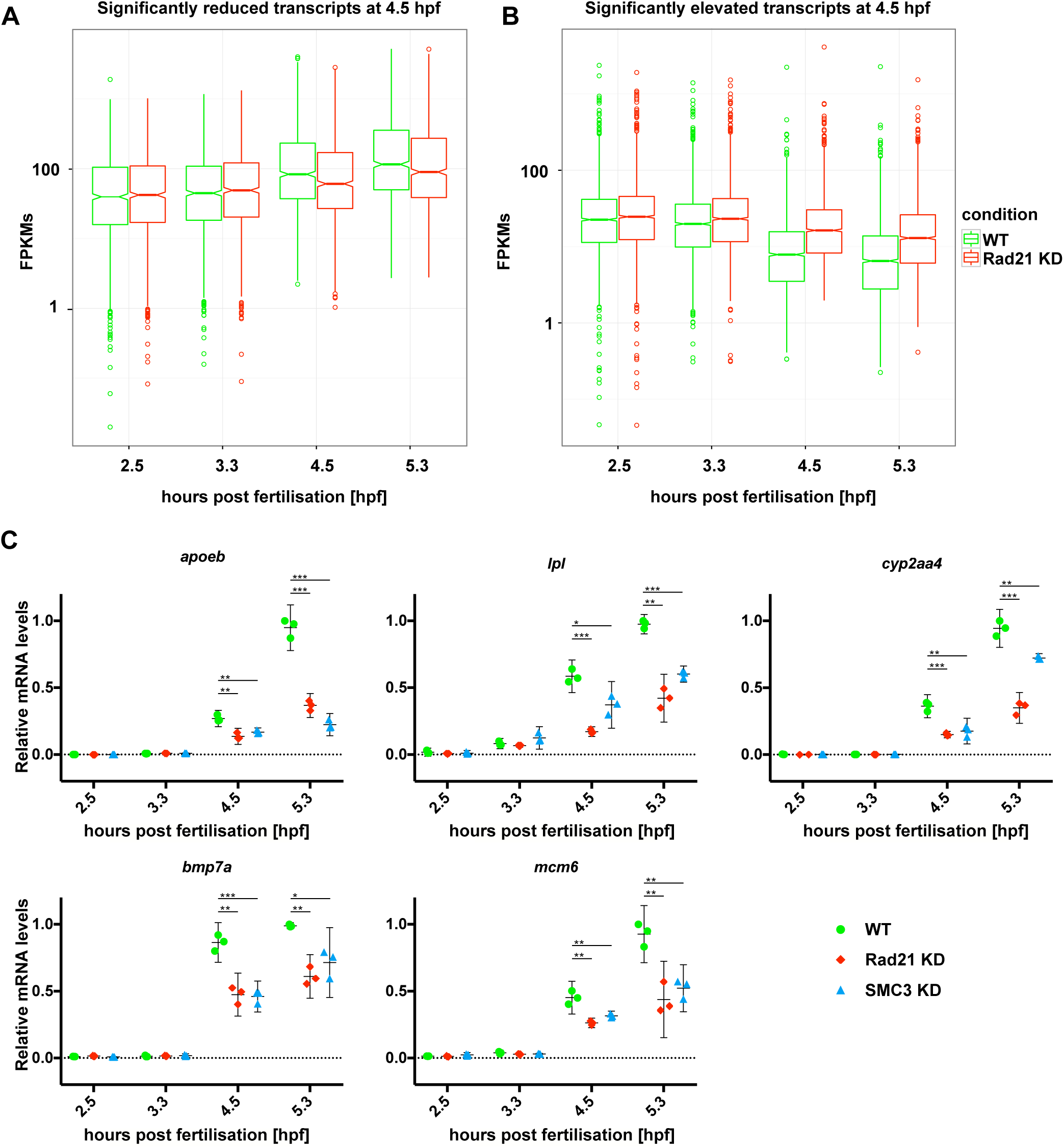
Cohesin depletion delays expression of zygotic genes. (A, B) Distribution of significantly differentially represented transcripts (FDR 0.05) in Rad21-depleted embryos over developmental time points (2.5-5.3 hpf). The bottom and top of the boxes represent the first and third quartiles, and the line within represents the median; notches represent confidence intervals. The whiskers denote the interval within 1.5 times the interquartile range (IQR) from the median. (A) FPKM over developmental time of 1,286 transcripts that were reduced in Rad21-depleted (KD) embryos. (B) FPKM over developmental time of 2,381 transcripts with elevated levels in Rad21-depleted (KD) embryos. (C) Quantitative RT-PCR of selected zygotically-expressed transcripts that were differentially represented in RNA-seq data. Embryos were injected at the one-cell stage with 1 pmol Rad21 or Smc3 morpholino respectively, and ~50 per condition were pooled for RNA extraction. Data were normalized to mitochondrial transcript *nd3* and shown as a scatter plot with means and 95% confidence intervals (3 biological replicates per condition). *p-*values (* <0.05, ** <0.01, *** <0.001 unpaired *t*-test).

While transcripts that are differentially represented upon Rad21 depletion were assignable to functional pathways (Fig. 4), transcripts responding to CTCF depletion were not. At the 4.5 hpf ‘dome’ stage, transcripts under-represented upon Rad21 depletion are involved in ribosome assembly, translation and RNA metabolism functions (Fig. 4A, Table S2). Over-represented transcripts reflect the maternal RNA landscape and are involved in energy systems and mitochondrial functions (Fig. 4B, Table S2).

**Figure 4:**
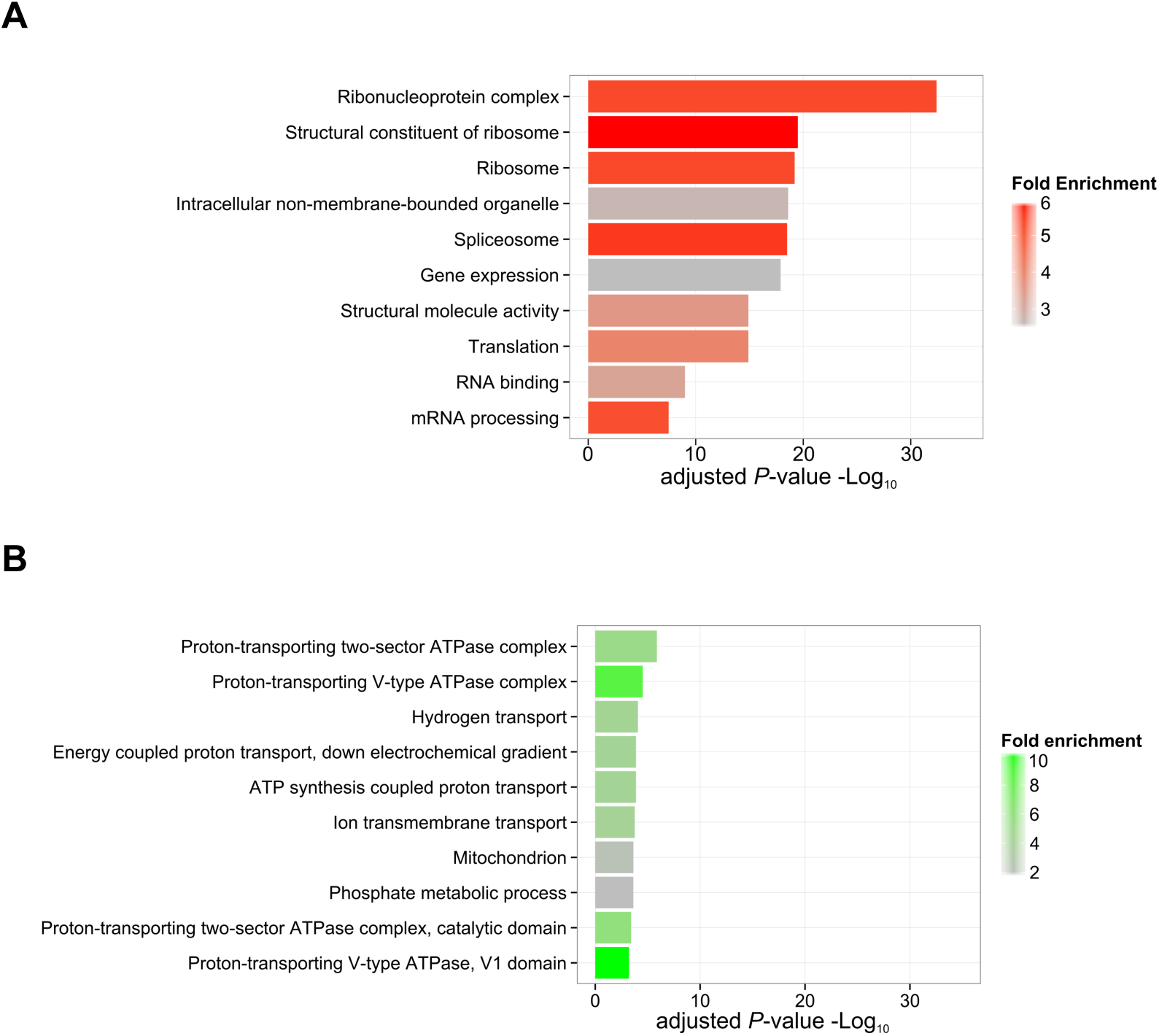
Gene ontologies of differentially represented transcripts in Rad21-depleted embryos. Enriched gene ontology (GO) terms and their binomial *p*-values with fold enrichment over expected number was derived using R package clusterProfiler to analyze differentially represented transcripts upon Rad21 depletion. (A) transcripts under-represented in Rad21-depleted embryos at 4.5 hpf; (B) transcripts over-represented in Rad21-depleted embryos at 4.5 hpf. The full list of GO terms that were enriched can be found in Table S2.

The RNA-seq data indicate that Rad21 (but not CTCF) depletion led to a delay in degradation of maternal mRNAs in combination with a delay in activation of zygotic genes, when compared to stage-matched embryos of equivalent morphology and cell number to wild type. Overall, our data suggests that cohesin is necessary for the timely transition to, and promotion of, maternal to zygotic transcription programs.

### Rad21 binding redistributes through ZGA

Considering the importance of Rad21/cohesin for progression to the zygotic transcription program (Figs 2-4), we decided to further investigate Rad21 function during ZGA. To determine the distribution of Rad21 on chromosomes in early development, we conducted chromatin immunoprecipitation followed by high throughput sequencing (ChIP-seq) in wild type embryos at 2.5, 4.5 and 10 hpf with custom antibodies against zebrafish Rad21 (Rhodes et al., 2010) (Fig. S5). At 2.5 hpf pre-ZGA, 2,011 enriched Rad21 peaks were detected on chromosomes. After ZGA, there was significant recruitment of Rad21 to chromosomes that increased over developmental time (Fig. 5A). By the 4.5 hpf ‘dome’ stage, wild type embryos had accumulated 7,144 significant Rad21 binding peaks, and by 10 hpf, there were 18,075 peaks in total. During ZGA and early development, Rad21 peak distribution shifts closer to the transcription start sites (TSS) of genes (Fig. 5A). The data suggest that pre-ZGA, Rad21/cohesin binds to few loci and is mostly excluded from genes, whereas post-ZGA, Rad21 binding accumulates at gene-dense regions. We performed ATAC-seq (Buenrostro et al., 2015) on wild type embryos at 2.5 hpf to determine if Rad21 binds to open chromatin regions at pre-ZGA. About half the accessible chromatin sites at 2.5 hpf also recruit Rad21 (Fisher’s exact test, right tail: *p* ≤ 1.00^−20^) (Fig. 5B), indicating that a subset of cohesin binding sites are located in the very few regions, 337 in total, which are accessible pre-ZGA. About 90% of the overlapping accessible regions correspond with satellite DNAs (Table S3; specific examples are shown in Fig. S6)

The remarkable redistribution of Rad21 binding to genes post-ZGA is exemplified by its recruitment at chromosome 4 (Fig. 5C). The long arm of chromosome 4 is gene-poor, has extensive heterochromatin, and replicates late. High densities of 5S ribosomal DNA (rDNA), small nuclear RNAs (snRNAs), half of all tRNAs, and 30% of all zinc finger domain genes are present on the long arm of chromosome 4 (Howe et al. 2013). Prior to ZGA, many of these loci were enriched for Rad21 (right side), whereas the RNAPII gene-rich region of chromosome 4 (left side) excluded Rad21 binding (Fig. 5C). Post-ZGA, Rad21 binding became increasingly enriched at the RNAPII gene-rich region of chromosome 4, and some of its pre-ZGA binding sites were lost (Fig. 5C).

**Figure 5:**
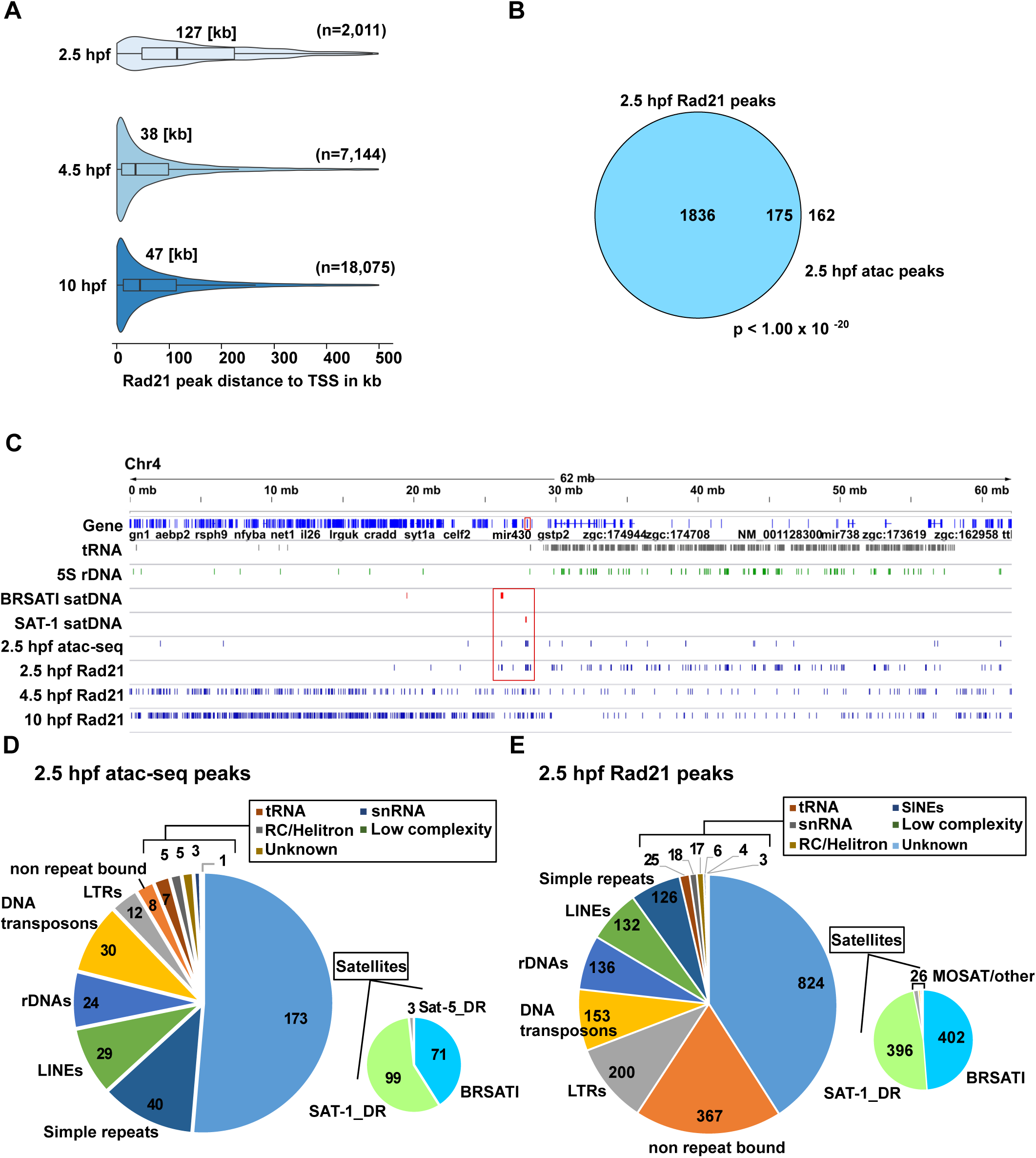
Rad21 binding redistributes to coding regions during ZGA. (A) Violin plots of the absolute distance to the TSS of Rad21-associated genes for stages 2.5, 4.5 and 10 hpf, including median values. The bottom and top of the boxes are the first and third quartiles, and the line within represents the median. The whiskers denote the interval within 1.5 times the interquartile range (IQR) from the median. (B) Overlap of ATAC-seq peaks at 2.5 hpf with Rad21 distribution at 2.5 hpf. The *p*-value was calculated using Fisher’s exact test (right-tail). (C) IGV genome browser view of chromosome 4. Rad21 locates to gene-poor regions on the long arm pre-ZGA but binds gene-rich regions on the short arm post-ZGA. Repetitive elements such as tRNAs, ribosomal RNAs (5S RNA) are highly enriched on chromosome 4 and overlap with Rad21. Satellite repeats (BRSATI and SAT-1) enriched at pericentromeric regions are also bound by Rad21 at 2.5 hpf. (red box). (D-E) Pie plots representing various repetitive elements overlapping pre-ZGA ATAC-seq peaks at 2.5 hpf and Rad21 peaks only at 2.5 hpf (E). 51% (173/337) of the ATAC-seq peaks and 41% (824/2011) of the Rad21 peaks associate with satellite repeats BRSATI and SAT-1.

Genes that recruit Rad21 pre-ZGA (within a 20 kb window) included *hsp70l*, *sox2*, *gata2a* and the *miR-430* complex (Fig. S7); and multiple zinc finger domain encoding proteins located on chromosome 4 (Fig. S8). Transcripts from *miR-430* and zinc finger domain encoding genes are expressed as early as the 64-cell stage, prior to the main wave of zygotic transcription (Heyn et al. 2014). The mature *miR-430* microRNAs mark a substantial amount of maternally deposited transcripts for degradation (Giraldez 2006). Interestingly, many zinc finger encoding genes marked by Rad21 binding are not expressed until post-ZGA (Fig. S8).

Besides association with regions on the long arm of chromosome 4, 41% (824/2,011) of the Rad21 peaks were found at satellite elements (satDNAs) located at pericentromeric regions of the genome (Figs 5D,E; S6, Table 1). Further classification of these elements shows that BRSATI and SAT-1 were among the highest enriched members of satDNAs found at Rad21 sites pre-ZGA representing over 70% of satDNAs identified. satDNAs represent less than 1% of the genome in zebrafish and are therefore significantly enriched (*p* value < 1.0^−20^, Fisher’s exact test) in the 2.5 hpf Rad21 peaks, whereas DNA transposons are relatively abundant accounting for 33% of the genome and are not significantly enriched. Long terminal repeats (LTRs) and rRNAs are also significantly enriched in pre ZGA Rad21 peaks (Table 1).

**Table 1:** Genome wide distribution of selected repeat elements and overlap with pre-ZGA Rad21 binding.

| **Repeat element** | **Number of elements in zv9** | **Rad21 overlap** | **p-values for fisher's exact test (right-tail)** | **Odds ratio** | **Genome wide coverage** |
| --- | --- | --- | --- | --- | --- |
| **LTRs** | 153185 | 200 | 7.15E-18 | 2.013 | 4.97% |
| **DNA transposons** | 1980516 | 153 | 1 | 0.143 | 33.88% |
| **rDNAs** | 3628 | 136 | 1.00E-20 | 204.286 | 0.04% |
| **BRSAT1** | 235 | 402 | 1.00E-20 | inf | 0.10% |
| **SAT-1 DR** | 176 | 396 | 1.00E-20 | inf | 0.06% |
| **MOSAT DR** | 10546 | 13 | 0.01641 | 2.026 | 0.29% |

### Rad21 locates to genes upon genome activation

After ZGA, there was significant recruitment of Rad21 to chromosomes that increased over developmental time (Fig. 5A). By the 4.5 hpf ‘dome’ stage, wild type embryos had accumulated 7,144 significant Rad21 binding peaks including ~3,000 that were gene-associated, and by 10 hpf, there were 18,075 peaks in total with 5,937 gene-associated (Figs 5A, 6A, Table S3). Rad21 binding was significantly over-represented in coding regions after ZGA. Furthermore, Rad21 binding was particularly enriched at promoters and 5′ untranslated regions (5′ UTR), as well as to exons, transcription termination sites (TTS), and 3′ UTRs (Fig. 6A).

**Figure 6:**
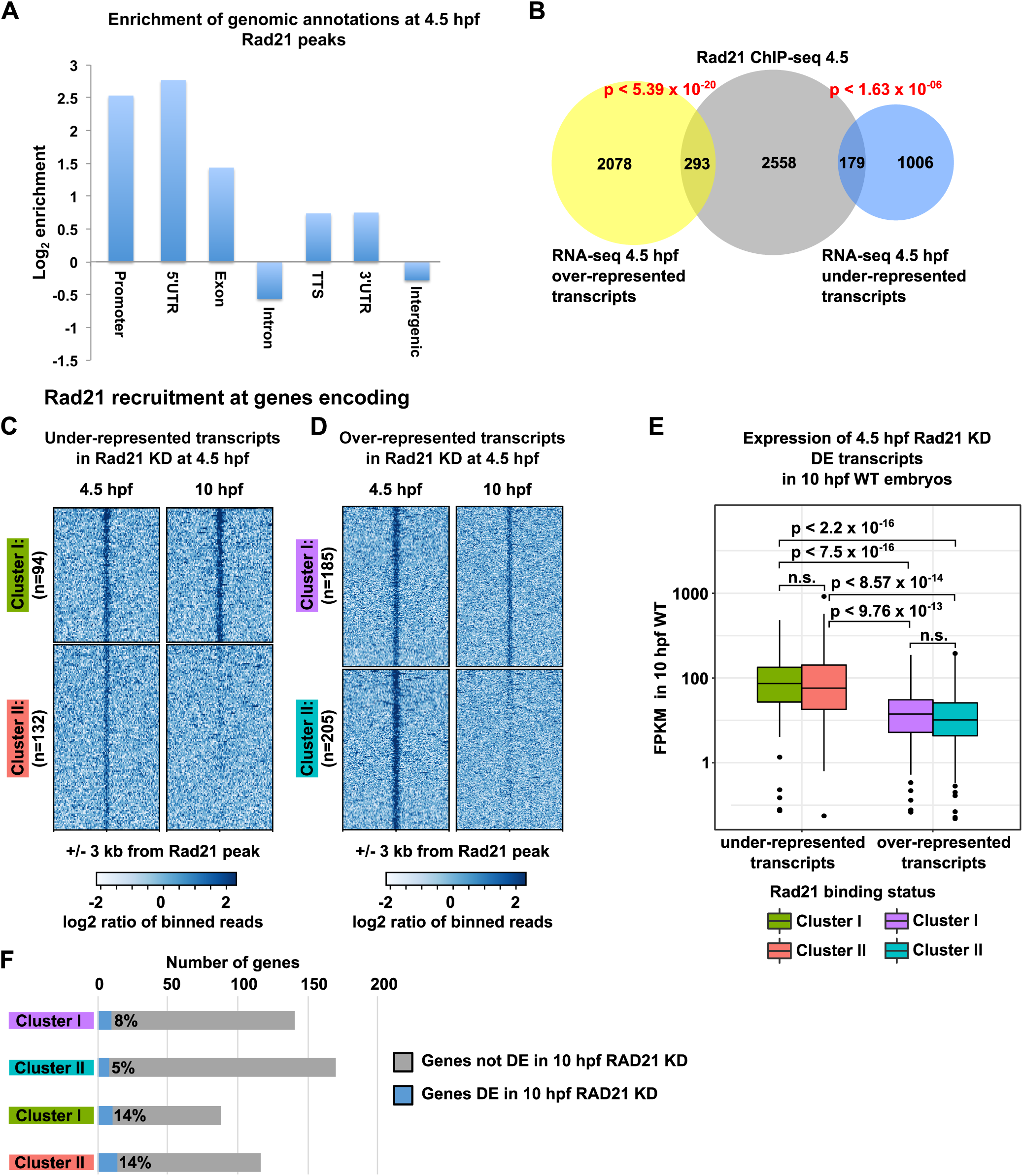
Post-ZGA Rad21 binding is enriched at genes and overlaps with differentially represented transcripts. (A) Enrichment of genomic features (3′ UTR, TSS, Exon, Intron, Promoter, 5′ UTR) at Rad21 binding sites. (B) Overlap between Rad21-bound genes and differentially represented transcripts upon Rad21 depletion was significantly enriched at 4.5 hpf (Fisher’s exact test: *p* ≤ 5.39^−20^, downregulated transcripts; *p* ≤ 1.630^−6^, upregulated transcripts). (C, D) Heat maps showing binding profiles of Rad21 at 4.5 hpf and 10 hpf, at regions associated with genes encoding over-represented transcripts (C) and under-represented transcripts (D) in 4.5 hpf Rad21-depleted embryos (KD). (E) Expression levels of genes associated with regions in (C) and (D) in 10 hpf wild type embryos. The bottom and top of the boxes are the first and third quartiles, and the line within represents the median. The whiskers denote the interval within 1.5 times the interquartile range. Rad21-bound genes with under-represented transcript levels upon Rad21 depletion (KD) at 4.5 hpf have higher FPKMs in 10 hpf wild type (WT) embryos than Rad21-bound genes with over-represented transcript levels upon Rad21 depletion at 4.5 hpf. All *p-*values were calculated using the Mann-Whitney-Wilcoxon test. (F) Number of differentially expressed (DE) genes in 10 hpf Rad21-depleted (KD) embryos.

Overall, 12% (293/2371) of over-represented transcripts, and 15% (179/1185) of under-represented transcripts were derived from genes that recruited Rad21 (Fig. 6B), implying the corresponding genes could be directly regulated by Rad21. The association of differential expression with bound genes is significant (Fig. 6B), even though relatively few dysregulated genes are bound. We used *k*-means clustering (*k*=2) to visualize Rad21 binding profiles over two subsequent developmental stages (4.5 and 10 hpf). About half of the regulated genes that contained Rad21 binding (58% for under-represented transcripts and 52% for over-represented) had lost that binding by 10 hpf (Fig. 6C,D), indicating that Rad21 is likely to be specifically associated with those genes during ZGA, and potentially involved in their direct regulation at that time. Following Rad21 depletion, when compared with over-represented transcripts, genes with under-represented transcripts at 4.5 hpf had higher transcription levels in wild type embryos by 10 hpf, irrespective of Rad21 binding (Fig. 6E). Only a small fraction (5-8%) of genes with over-represented transcripts at 4.5 hpf also showed altered expression at 10 hpf. Genes found to be downregulated at 4.5 hpf were more likely to also be differentially expressed at 10 hpf (14%), but there was no difference between genes that gain or lose Rad21 binding at 10 hpf (Fig. 6F, clusters I and II, respectively). This indicates that a small subset of genes bound by Rad21 during ZGA are affected later in development by Rad21 depletion. However, although it is likely that some of the bound genes may be regulated directly, a larger fraction appears to be regulated indirectly.

Our results indicate that Rad21 is present at repetitive sequences and ncRNA genes prior to ZGA, with a transition to RNAPII genes at ZGA, once transcription starts. The marked enrichment of Rad21 at genes through developmental time suggests that cohesin may facilitate their expression. However, because many more genes are regulated by Rad21 depletion than are bound by Rad21, it is unlikely that direct gene regulation by cohesin explains the delay in ZGA.

### Rad21 binding coincides with active histone marks and sites occupied by pluripotency factors Pou5f3 and Sox2

To further investigate a possible role for Rad21 in ZGA, we sought to determine if Rad21 binding coincides with other hallmarks of gene activation, including H3K4me1 and H3K27ac enhancer modifications, H3K4me3 marks associated with active gene promoters, and H3K27me3 modification of polycomb-repressed genes (Vastenhouw and Schier, 2012). For this analysis, we surveyed defined regions centered on Rad21 binding sites for enrichment of these modified histones by comparing Rad21 ChIP-seq data to publically available histone ChIP-seq data (Bogdanovic et al., 2012; Zhang et al., 2014b) at peri-ZGA time points. (Fig. 7A). 48% of the Rad21 peaks (3,416/7,144) overlapped with at least one of the enhancer and promoter associated marks, H3K4me1, H3K4me3 and H3K27ac (hypergeometric test, *p* ≤ 2.72^−2803^) (Fig. 7A, Table S4). A smaller set of 244 peaks was significantly associated with H3K27me3 (hypergeometric test, *p* ≤ 2.72^−34^). Less than half of overlapping peaks were at TSSs (Table S4). Therefore, cohesin binding significantly coincides with histone marks that are associated with active chromatin at ZGA.

**Figure 7:**
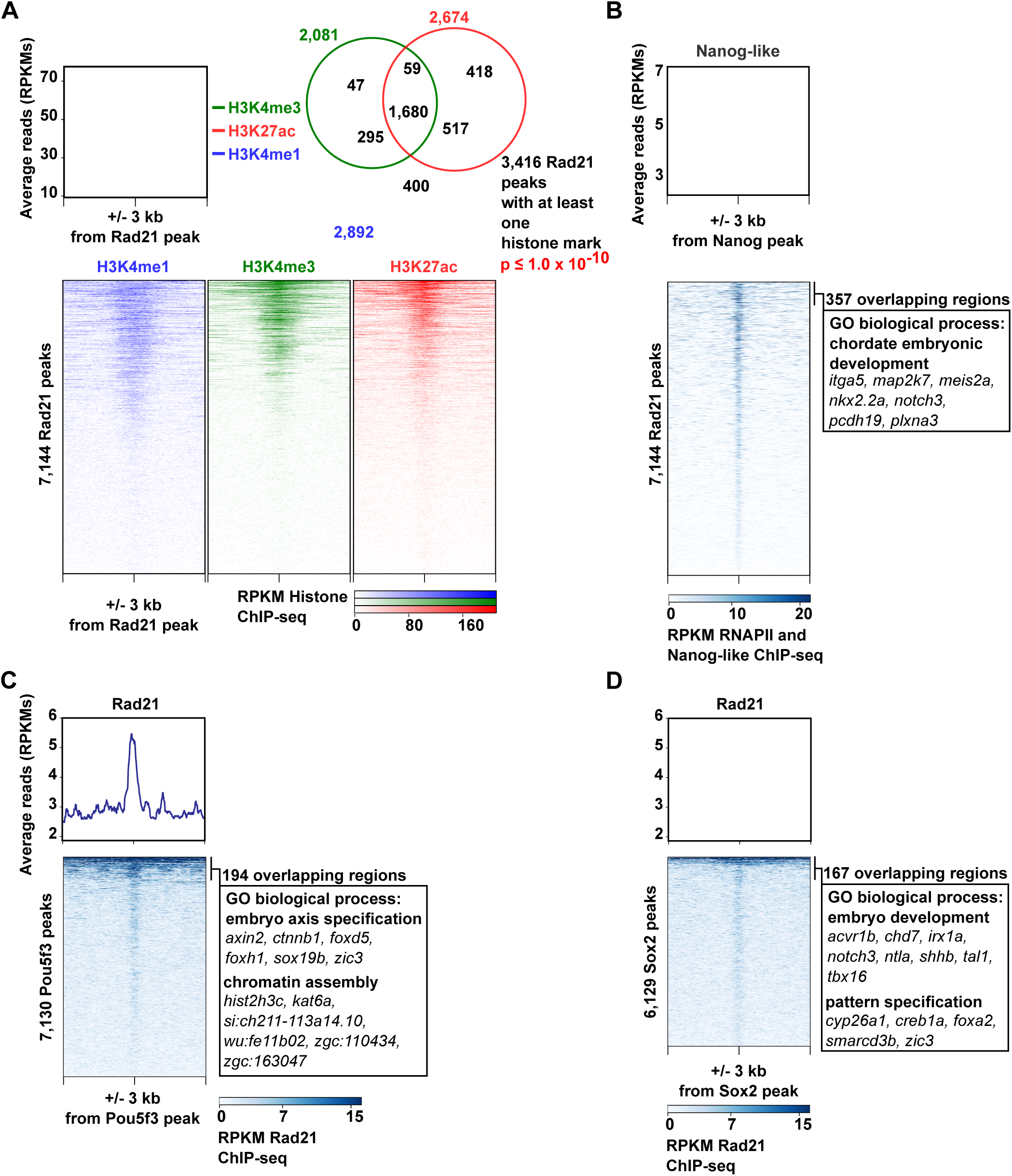
A subset of Rad21 binding sites coincide with occupancy of active histone marks and pluripotency factors, Nanog-like, Pou5f3 and Sox2. (A) Histone modifications at Rad21 binding sites. Heat maps and average profiles showing enrichment of histone marks over defined regions centered on individual Rad21 peaks at 4.5 hpf. Heat maps are ordered by decreasing enrichment for each histone modification independently. Weighted Venn diagram of Rad21 peaks overlapping with different histone modification peaks from 4.5 hpf embryos. For Rad21 overlap with histone marks, a hypergeometric test was used. (B) Heat maps and average profiles showing enrichment of Nanog-like binding at Rad21 peaks at 4.5 hpf. Significantly enriched gene ontologies of Rad21 and Nanog-like overlapping regions. (C) Heat maps and average profiles showing Rad21 enrichment at Pou5f3 peaks. Significantly enriched gene ontologies of Rad21 and Pou5f3 overlapping regions. (D) Heat maps and average profiles showing Rad21 enrichment at Sox2 peaks. Significantly enriched gene ontologies of Rad21- and Sox2-overlapping regions.

The transcription factors Nanog-like, Pou5f3 and the SoxB1 family (including sox2, sox3, sox19a and sox19b) are homologs of mammalian pluripotency factors, and are thought to act as pioneering factors in zebrafish ZGA (Lee et al., 2013; Leichsenring et al., 2013). Because Rad21 depletion delayed ZGA (Fig. 2), we were interested to know whether Rad21/cohesin binding coincides with genomic locations of these activators of early gene expression. Publically available ChIP-seq data for Nanog-like (C. Xu et al. 2012), Pou5f3 and Sox2 (Leichsenring et al., 2013) was obtained (Table S4) and compared to Rad21 binding at 4.5 hpf. There was a small but significant overlap between Rad21 and pluripotency factor binding sites (Table S4 and Fig. 7B-D). Regions with overlap of Rad21 and pluripotency factors were enriched for developmental, chromatin assembly, and pattern specification ontologies (Fig. 7B-D). The coincidence of a subset of cohesin binding with these known transcriptional activators suggests that cohesin may be involved in regulating selected Sox2 and Pou5f3 targets.

### Rad21 depletion disrupts nuclear structure and RNA Polymerase II clustering at ZGA

The combined data above point to a generalized role for Rad21 in early zygotic transcription. Given the known role of cohesin in the local spatial organization of chromatin (Merkenschlager and Nora, 2016), we addressed the possibility that Rad21/cohesin might contribute to ZGA through global organization of chromatin architecture.

We used antibodies to Nucleolin and RNAPII to visualize nucleoli and RNAPII clustering, respectively, immediately post-ZGA. Immunofluorescence analysis in 4.5 hpf ‘dome’ stage embryos revealed the formation of nucleoli (Fig. 8A,A′) and discrete RNAPII clusters that may represent transcription foci (Fig. 8D,D′). Strikingly, depletion of Rad21 severely and significantly disrupted the formation of nucleoli (Fig. 8B,B′,C - *p* ≤ 4.7^−14^) and RNAPII clusters (Fig. 8E,E′,F - *p* ≤ 8.8^−7^) at this developmental stage, with these markers exhibiting a more fragmented appearance.

**Figure 8:**
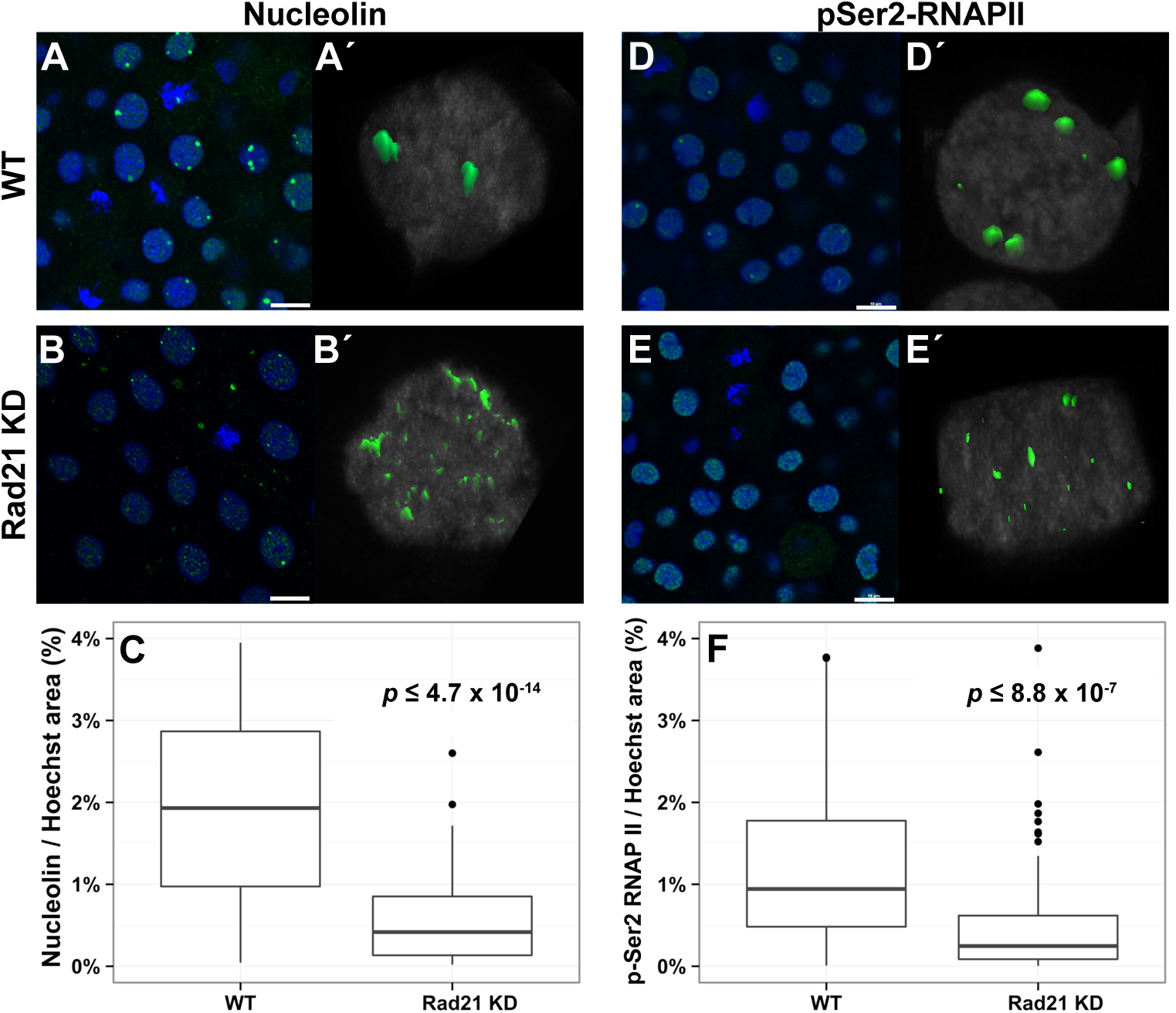
Formation of sub-nuclear structures in post-ZGA embryos is compromised by Rad21 depletion. Rad21-depleted (Rad21 KD) and wild type (WT) stage-matched control embryos at 4.5 hpf were fixed and stained with the indicated antibodies. For all images, nuclei were counterstained with Hoescht, and the scale bar is 10 μm. (A, A′) Nucleolin staining (green) in wild type is shown in a field of cells (A) and in a z-stack maximum projection of a single representative nucleus (A′), and indicates the presence of normal nucleoli. (B, B′) Nucleolin staining (green) in Rad21-depleted embryos is shown in a field of cells (B) and in a z-stack maximum projection (B′), and indicates nucleolar dispersion following abrogation of Rad21. (C) Quantification of the area of Nucleolin relative to the size of the nucleus in stage-matched wild type embryos compared with Rad21-depleted embryos at 4.5 hpf (see methods; n=6 for both conditions) shows that nucleoli fragmentation is significant. Around 200 nuclei for each condition were imaged and analyzed. (D, D′) Staining for the elongating form of RNA polymerase II (p-Ser2-RNAPII) (green) in wild type is shown in a field of cells (D) and in a z-stack maximum projection of a single representative nucleus (D′), and indicates clustering of RNAPII into foci. (E, E′) p-Ser2-RNAPII staining (green) in Rad21-depleted 4.5 hpf embryos is shown in a field of cells (E) and in a z-stack maximum projection (E′) shows disruption of RNAPII foci upon Rad21 depletion. (F) Quantification of RNAPII foci relative to the size of the nucleus shows statistically significant disruption of RNAPII clustering in Rad21-depleted embryos compared with controls (see methods; n=6 for both conditions). Around 200 nuclei for each condition were imaged and analyzed. All *p* values were calculated by applying a *t*-test with an unpaired fit and assuming a parametric distribution.

Ribosomal DNA (rDNA) is contained within the nucleolus and active rDNA interacts with Nucleolin (Cong et al. 2012a). In support of a direct role for Rad21 in nucleolar organization, we found that Rad21 is enriched near 5S ribosomal DNA repeats (Cong et al. 2012b) on chromosomes 4 (Fig. 5C), 18 and 22 (Table S3).

Thus, depletion of Rad21 dramatically affects nuclear organization by the time of ZGA. Our results further suggest that Rad21 recruitment to genes at this crucial developmental stage influences the formation of RNAPII foci that could represent early sites of transcription. Transcripts from many genes are affected by Rad21 depletion in a manner consistent with an overall delay in ZGA (Figs 2, 3), although few of these genes are directly bound by Rad21 (Fig. 6). Global disruption of chromosome organization by Rad21 depletion provides a possible mechanism for this observation.

## Discussion

Altogether, our results point to a global role for Rad21/cohesin in facilitating ZGA in zebrafish embryos. Rad21 locates to active regions of the genome, including genes expressed at ZGA, while Rad21 depletion interferes with gene expression. Rad21 depletion also affects nuclear integrity and RNAPII clusters, raising the possibility that cohesin plays a role in organizing a chromatin structure that is permissive for transcription around the time of ZGA.

### Transcriptional changes at ZGA following Rad21 or CTCF depletion

Owing to their important combinatorial roles in genome organization (Vietri Rudan and Hadjur, 2015), we expected Rad21/cohesin and CTCF depletion to have similar effects on activation of the zygotic genome. We were surprised to find that, upon depletion, their effects were quite different. We previously showed that a modest depletion of Rad21 by morpholino (Schuster et al., 2015) or mutation (Horsfield et al., 2007) can have striking effects on the transcription of specific genes. By contrast, CTCF had to be dramatically depleted to affect transcription, and doing so resulted in high levels of mortality (data not shown and Marsman et al., 2014). CTCF is essential to the integrity of the nucleus; our data suggest that a small amount of CTCF may be sufficient for this function, and that CTCF depletion beyond this level is lethal in embryos in which cells proliferate rapidly. Consistent with this, maternal and zygotic depletion of CTCF leads to apoptosis and is lethal at preimplantation stages in mice (Moore et al., 2012; Wan et al., 2008). In contrast to CTCF, partial Rad21 depletion generated multiple robust biological effects at zebrafish ZGA, and this tractability encouraged us to focus our study on Rad21/cohesin.

### Rad21 depletion delayed ZGA and dysregulated transcripts in distinct pathways

Rad21 depletion dramatically altered the transcript complement in embryos just post-ZGA, reflecting an overall delay in transition from the maternal to zygotic transcription program. Among the top downregulated gene ontology categories were ribosome assembly, RNA processing and translation. These processes are also compromised by disrupting cohesin function in yeast and mammalian cells (Bose et al., 2012). Xu *et al.* previously demonstrated that translational defects in zebrafish and mammalian cell cohesin mutants were chemically rescued by L-Leucine stimulation of the TOR pathway (Xu et al., 2016; Xu et al., 2015). Our data are consistent with these observations that translational mechanisms require normal cohesin function. Moreover, cell proliferation dominates early zebrafish development and requires high levels of translation, consistent with the emergence of these biological pathways as the most significantly affected by cohesin depletion just post-ZGA. Compromising this aspect of the normal gene expression program will almost certainly affect embryogenesis, consistent with mutations in cohesin causing the human developmental disorder, Cornelia de Lange Syndrome (CdLS).

Although cell proliferation is central to early development, in Drosophila and zebrafish, ZGA is independent of, or upstream of cell cycle number and checkpoint regulators (Blythe and Wieschaus, 2015; Zhang et al., 2014a; Zhang et al., 2017). Consistent with these observations, we found that delay in ZGA occurred in Rad21-depleted embryos that had the same number of cells as controls. However in zebrafish, ZGA does reflect replication timing in the early embryo (Siefert et al., 2017), and we cannot rule out the possibility that replication timing is affected in our experiments.

### Cohesin binding is restricted to select transcript-encoding locations pre-ZGA

Rad21 was generally excluded from genes pre-ZGA with some notable exceptions. Genes that recruited Rad21 pre-ZGA included *hsp70l*, *sox2*, *gata2a*, and the *miR-430* complex. Of these, *gata2a* and *miR-430* are expressed pre-ZGA (Heyn et al., 2014), and cohesin binding to these locations was reduced once embryos transited through ZGA. Interestingly, miR-430 is responsible for targeting maternal transcripts for clearance (Bazzini et al., 2012; Giraldez et al., 2006), and this raises the possibility that a proportion of maternal transcripts with delayed degradation upon Rad21 depletion could be accounted for by dysregulated *miR- 430*.

Other genes that recruit Rad21 pre-ZGA are generally not expressed at that time. For example, *sox2* mRNA is maternally provided, and is involved in transcription of early-expressed zygotic genes in zebrafish (Lee et al., 2013). The zygotic *sox2* gene is expressed post-ZGA (Heyn et al., 2014). Significantly, the timing of *sox2* expression post-ZGA coincides with a redistribution of Rad21 peaks at the *sox2* gene. In addition, Rad21 is recruited to several zinc finger protein-encoding genes pre-ZGA that are expressed at post-ZGA stages of development. Recruitment of Rad21 to silent loci pre-ZGA could indicate that cohesin has a function there, perhaps to mark their later expression.

### Cohesin is enriched at pericentromeric satellite DNA repeats

Prior to ZGA, cohesin is highly enriched at satellite DNAs found at pericentromeric regions, which represent less than 1% of the genome, as well as ncRNA genes. Various satellite sequences in somatic cells are packaged into constitutive heterochromatin, which is characterized by high compaction, enrichment of repressive histone modifications, transcriptional quiescence, and late replication. Most of these attributes are absent in pre-ZGA embryos, and the satellite sequences seem to take on these features successively as the embryo develops (Borsos and Torres-Padilla 2016). ATAC-seq indicates that the satellite DNAs are highly accessible pre-ZGA. It is possible that cohesin is sequestered there merely because this chromatin is open, and thus satellite DNA serves to keep cohesin away from RNAPII genes prior to genome activation. Cohesin depletion results in organizational changes of nucleoli, which could interfere with satellite-dependent heterochromatin formation at ZGA.

### How does cohesin contribute to transcription of the zygotic genome?

At post-ZGA stages, thousands of genomic locations recruit Rad21, and markedly include genic features such as promoters, TSSs, termination sites, 3′ and 5′ UTRs and exons. Subsequently, Rad21 is increasingly enriched TSSs, notably at sites co-enriched in histone modifications indicative of active promoters and enhancers (Vastenhouw and Schier, 2012; Vastenhouw et al., 2010). Gene-associated Rad21 significantly overlapped with occupancy of the pluripotency factors Pou5f3 and Sox2 at similar time points. This raises the possibility that the pluripotency factors pioneer sites of zygotic transcription (Lee et al., 2013) and recruit cohesin at a subset of these to keep these regions in an ‘open’ configuration. In support of this idea, nucleosome density increases following cohesin loss (Yan et al., 2013), suggesting that cohesin acts to keep chromatin open. Moreover, nucleosome organization is a key feature of ZGA; in zebrafish, nucleosomes are strongly positioned at promoters (Zhang et al., 2014b) at a stage that is coincident with cohesin binding.

Cohesin may also operate at a nuclear structural level to regulate ZGA. Consistent with previous observations in yeast and human cells (Bose et al., 2012; Harris et al., 2014), Rad21/cohesin is essential for the formation of nucleoli in post-ZGA zebrafish embryos. Loss of nucleoli could have global effects on zygotic transcription and translation, as was observed in this study and others (Xu et al., 2015). In addition, reduction in Rad21 just post-ZGA resulted in dispersion of RNAPII foci that could represent transcription factories. Loss of chromosome architecture owing to cohesin depletion could lead to an inability to assemble a transcription-competent genome structure.

A combination of the factors described above could lead to global dysregulation of the zygotic transcription program, and these factors are indicative of roles for cohesin at multiple levels at ZGA (Fig. 9).

**Figure 9:**
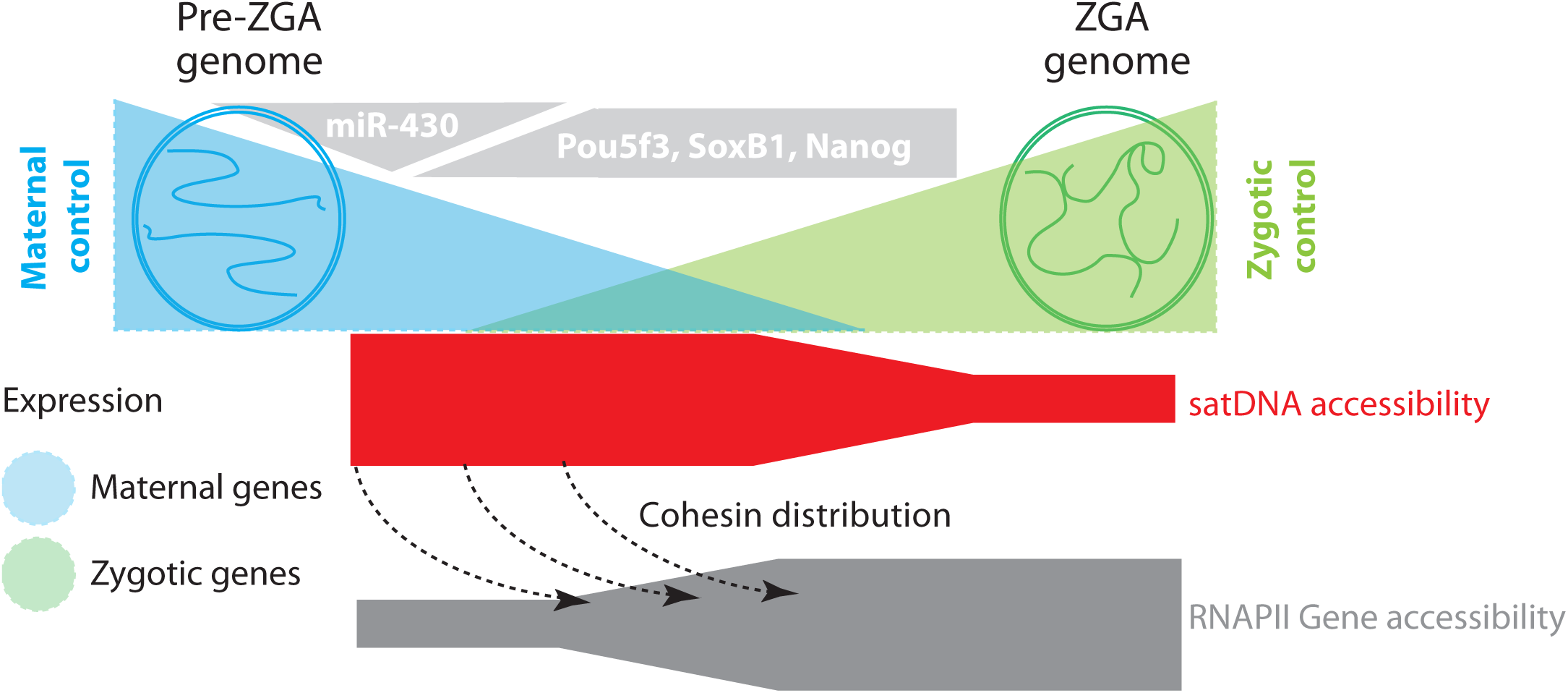
Model of potential mechanisms for cohesin regulation at ZGA. Up until ZGA, cohesin locates to accessible regions of the genome, including miR-430 and satellite DNA. As embryos transit through ZGA, cohesin relocates to RNAPII genes. Access of cohesin to zygotic genes may be regulated by transcription factors Pou5f3, SoxB1 and Nanog. Enrichment of cohesin at genes may contribute to forming transcription competent local and global chromatin structures.

A spectrum of multifactorial human developmental disorders known as the ‘cohesinopathies’ arise from mutations in cohesin regulators or cohesin subunits (Ball et al., 2014; Bose and Gerton, 2010; Horsfield et al., 2012; Skibbens et al., 2013). Our study raises the possibility that germline cohesinopathy mutations could lead to global alteration of the zygotic transcription program right from the start of development, perhaps explaining the diversity of phenotypes observed in cohesinopathy patients.

## Materials and Methods

### Zebrafish and microinjection

Zebrafish were maintained under standard conditions (Westerfield, 1995). The University of Otago Animal Ethics Committee approved all zebrafish research. Morpholino oligonucleotides (MOs) were obtained from GeneTools LLC and diluted in water. MO sequences were Rad21 5′-AGGACGAAGTGGGCGTAAAACATTG-3′; and CTCF 5′-CATGGGTAATACCTACATTGGTTAA-3′ (targeting the ATG), 5′-CCAAAACAGATCACAAACCTGAAAG-3′ (targeting the splice site of intron 2); and Smc3 5′-TGTACATGGCGGTTTATGC-3′ (targeting the ATG) as described previously (Marsman et al., 2014; Rhodes et al., 2010; Schuster et al., 2015). For microinjection, 1 nl containing 1.0 pmol (for embryos up to MZT) or 0.25-0.5 pmol (for embryos grown post-MZT) of each MO was injected at the 1-cell stage. CTCF MOs were combined in an equimolar ratio. For mRNA rescue of the Rad21 MO, embryos were injected with MO from one needle and rescue mRNA (200 pg) from a second needle. Mutant *rad21*^*nz171*^ mRNA (Horsfield et al., 2007) was used a control.

### RNA extraction

Wild-type embryos were collected at the one-cell stage, synchronized and either morpholino-injected or kept as control and allowed to develop to the desired stage (2.5, 3.3, 4.5, 5.3, 10 hpf) at 28 °C. Three biological replicates each containing total RNA from 100 pooled embryos were isolated using the NucleoSpin^®^ RNAII Kit (Macherey-Nagel). The quality of the RNA was confirmed using the Agilent 2100 Bioanalyzer, all samples had RIN >9.

### RNA sequencing, read mapping and bioinformatics analysis

Triplicate RNA samples from morphologically stage-matched embryos were sequenced to compare expression profiles over time. Strand-specific libraries were prepared using the TruSeq stranded total RNA-ribozero kit (Illumina) and 100-bp paired-end sequencing was performed to depth of 10 million reads per library on an Illumina HiSeq 2000. On average, 19 million 100 bp paired-end reads per library were generated. These were then adapter and quality trimmed using cutadapt (Martin, 2011) and SolexaQA (Cox et al., 2010). Each sequencing data set was independently mapped to the zebrafish genome with a bowtie2 index generated from Danio_rerio.Zv9.70 (Ensembl) downloaded from Illumina’s iGenomes collection. Zebrafish genome danRer7[Zv9] was used to provide known transcript annotations from Ensembl using TopHat2 (version 2.0.9) (Kim et al., 2013) with the following options: “tophat2 --GTF genes.gtf --library-type fr-firststrand -p 24 --mate-inner-dist −8 -- mate-std-dev 6 zv9” (on average, 75.38% reads mapped uniquely to the genome). Transcriptomes were assembled with Cufflinks (version 2.2.0) (Trapnell et al., 2010) using options: ‘cufflinks -p 32 --GTF genes.gtf’ and differential expression analysis between control and knockdown embryos was performed using Cuffdiff. A FDR corrected *p*-value of 0.05 was applied as the cut-off to identify differentially regulated transcripts. The R package DESeq2 was used to compare expression profiles over time. The R package clusterProfiler (Yu et al., 2012) was used to identify enriched Gene ontology terms in up-regulated and down-regulated gene lists using a cut-off of 0.05 FDR corrected *p*-value.

### Quantitative PCR

From RNA-seq data, five candidate genes were selected for confirmation by quantitative PCR following Rad21 and Smc3 depletion. Embryos were collected at four stages (2.5, 3.3, 4.5 and 5.3 hpf), RNA extracted as above, and cDNA synthesized (qScript). Quantitative RT-PCR was performed with primers designed to each of the five candidates (Table S5). Primers were designed to span exon-exon junctions to amplify only processed mRNA transcripts. Expression was normalized to the mitochondrial gene *nd3* (Table S5).

### Antibodies

Anti-Rad21 (Rhodes et al., 2010) and anti-CTCF (Marsman et al., 2014) were raised in rabbit against a 15 amino acid peptide of each of the zebrafish proteins, GenScript Corporation, USA. Commercial primary antibodies were: mouse anti-γ-tubulin (T5326; Sigma-Aldrich), anti-nucleolin (ab22758), anti-RNA polymerase II CTD repeat YSPTSPS (phospho S2) (ab5095) (Abcam), anti-SMC3 (D47B5) rabbit mAb #5696 (Cell Signaling). Secondary antibodies were: goat anti-Rabbit IgG (H+L) (#A-11008, Thermo Fisher Scientific), IRDye^®^-conjugated antibodies (#926-68070 and #926-32211, LiCor).

### Immunoblot analysis

Following dechorionation and deyolking, zebrafish embryos were lysed in RIPA buffer and equal amounts of protein were separated by electrophoresis on 10% polyacrylamide gels. Proteins were transferred to nitrocellulose (Thermoscientific) and incubated with mouse anti-γ-tubulin (1:5000) and rabbit anti-rad21 (1:500), secondary antibodies were the IRDye^®^-conjugated antibodies (1:15,000). Blots were visualized with the Odyssey^®^ CLx Infrared imaging system (LiCor). Band intensities were quantified using Image Studio 4.0 Software (LiCor).

### Chromatin immunoprecipitation (ChIP) sequencing and analysis

Chromatin was prepared from two independent collections of pooled embryos (n=2000) for 2.5 hpf stage embryos and (n=1000) for 4.5 hpf and 10 hpf embryos as described in (Lindeman et al., 2009). Briefly, embryos were dechorionated using a syringe with a 21G needle, fixed in 1% formaldehyde 10 minutes at room temperature. Fixation was stopped by adding glycine to a final concentration to 0.125 M and incubation on ice for 5 minutes. Fixed embryos were then washed three times in ice cold 1x PBS, snap frozen and stored at −80°C until use. After cell lysis, chromatin was sheared to 200-500 base pairs using a S220 Focused-ultrasonicator (Covaris) with the following settings per cycle: peak power = 70, duty factor = 5, cycles of bursts = 200, time=30 s. Individual cycle numbers were optimized for each stage. Chromatin from pre-MZT embryos needed 6 cycles of sonication to reach the desired 200-500 bp range, whereas chromatin isolated from 4.5 hpf and 10 hpf stages required 10 cycles. Cell debris was removed by centrifugation. To provide standardized input for each ChIP experiment, chromatin was diluted to A_260_=0.25. For each ChIP, 6 µg of Rad21 antibody per 10 µl Dynabeads and 100 µl chromatin was incubated overnight at 4 °C. After elution, ChIP DNA and input controls were purified and precipitated with ethanol.

The ThruPLEX^®^ DNA-seq Kit (Rubicon Genomics, USA) was used to prepare the 2.5 hpf sample libraries for sequencing. 125-bp paired-end sequencing was performed to a depth of 20-50 million reads per library on an Illumina HiSeq 2500^TM^ by New Zealand Genomics Limited, NZ. Libraries for the 4.5 hpf and 10 hpf samples were constructed and sequenced at the Beijing Genomics Institute (China), yielding 20 million 50 bp single-end reads per sample. After adapter and quality trimmed using cutadapt (Martin, 2011) and SolexaQA (Cox et al., 2010), reads were aligned to the Zv9 genome assembly using bowtie2 (Langmead and Salzberg, 2012) (version 2.2.1.) with default settings. 85% of the raw reads could be aligned except for Rad21 ChIP samples from 2.5 hpf, which had a 40% mapping rate. As an alternative to bowtie2, we used the aligner SHRiMP2 (David et al., 2011), increasing the mapping rate to 79% for the 2.5 hpf Rad21 IP samples. Peak finding and downstream data analysis was performed using HOMER (Nagy et al., 2013) and MACS2 (Zhang et al., 2008). Peaks were defined at a 0.1% estimated false discovery rate. Repetitive elements were obtained from the Repeatmasker database (Tarailo-Graovac and Chen, 2009) and overlapped with Rad21 peaks using Bedtools2 (Quinlan and Hall, 2010). Heat maps were generated using the log_2_ ratios of the binned reads comparing ChIP input and IP samples using deepTools2 (Galaxy version 2.2.3.0) bamCompare (Galaxy version 2.2.3), computeMatrix (Galaxy version 2.2.5) and plotHeatmap (Galaxy version 2.2.5) (Ramirez et al., 2016).

### Preparation and sequencing of ATAC-seq libraries

The ATAC-seq libraries from zebrafish embryos were prepared as previously described (Buenrostro et al., 2015) with some modification. Embryos were collected at the 256-cell stage and dechorionated using pronase. Yolk was removed using deyolking buffer as previously described (Link et al., 2006). 75,000 cells were used to prepare the libraries. Libraries were pooled and sequenced on an Illumina HiSeq. Data was aligned to Zv9 using bowtie2 and peaks for each replicate were called using MACS2 (Zhang et al., 2008). Peaks identified in both replicates were used for downstream analysis.

### Flow Cytometry

Around 100 embryos at 4.5 hpf (WT and MO injected) were dechorionated and deyolked. Cells were fixes with 100% ethanol overnight. Fxcycle™ PI/RNase staining solution (Thermo Fisher Scientific) was used to stain DNA in cells. Flow cytometric acquisitions were performed on a FACSCALIBUR (BD). Analyses were performed using FlowJo software (Treestar).

### Whole mount immunofluorescence

Embryos were fixed, dehydrated, and stored in 100% methanol at −20 °C. For staining, embryos were rehydrated in methanol/PBT, incubated in 150 mM Tris-HCl pH 9, followed by heating at 70 °C for 15 min (Inoue and Wittbrodt, 2011). Embryos were washed and blocked in 5% sheep serum, 2 mg/ml BSA in PBT. Primary antibodies were used at 1:1000, and the secondary antibody at 1:1000 dilution together with 1:1000 Hoechst stain (Thermo Fisher Scientific) in blocking buffer. Embryos were washed in PBT and stored in DAKO mounting media until image acquisition. Confocal immunofluorescence images were acquired with a confocal microscope (Nikon C2, Nikon) using a CFI Plan Fluor NA 0.3/10× objective and a NA 1.4/60× oil immersion objective.

### Image quantitation

The Imaris software package with default parameters was used to quantify numbers of nuclei per embryo from z-stacks of Hoechst stained embryos. For quantifying nucleolar integrity and RNAPII foci, a single focal plane was obtained through the center of the nucleus. To quantify the area of immunodetected Nucleolin or RNAPII relative to the size of the nucleus, a constant threshold setting in NIS-elements imaging software (Nikon) was used for wild type and Rad21-depleted embryos. The area of the nucleus outlined by the Hoechst staining defined regions of interest for which the pixel area of Nucleolin or RNAPII was measured.

### Statistical analysis

Statistical tests were performed using R (R Foundation for Statistical Computing, 2015).

### Data deposition

All datasets can be found at the Gene Expression Omnibus (GEO) GSE84602.

## Acknowledgments

The authors would like to thank Dale Dorsett for advice on ChIP-seq analysis, and Noel Jhinku for expert management of the zebrafish facility. This research was funded by Royal Society of NZ Marsden Fund [grant numbers 11-UOO-027, 16-UOO-072 to JAH], and Gravida National Center for Growth and Development [Doctoral Scholarship to MM].

